# Mitofusin 1 is required for the oocyte-granulosa cell communication that regulates oogenesis

**DOI:** 10.1101/498642

**Authors:** Thiago S. Machado, Karen F. Carvalho, Bruna M. Garcia, Amanda F. Zangirolamo, Carolina H. Macabelli, Fabrícia H. C. Sugiyama, Mateus P. Grejo, J. Djaci Augusto Neto, Fernanda K. S. Ribeiro, Fabiana D. Sarapião, Flávio V. Meirelles, Francisco E. G. Guimarães, Lena Pernas, Marcelo M. Seneda, Marcos R. Chiaratti

## Abstract

Mitochondrial function, largely regulated by the dynamics of this organelle, is inextricably linked to oocyte health. While the proteins that modulate mitochondrial fusion, Mitofusin 1 (MFN1) and 2 (MFN2), are required for embryogenesis, their role in oocyte development remains unclear. Here we show that the oocyte-specific deletion of *Mfn1*, but not *Mfn2*, results in a complete loss of oocyte growth and ovulation due to a block in folliculogenesis at the preantral-to-antral follicle transition. We pinpoint the loss of oocyte ovulation to disrupted oocyte-somatic cell communication – *Mfn1*-null oocytes are deficient for the production of the important somatic cell signaling factor GDF9. Unexpectedly, the double loss of *Mfn1* and *Mfn2* mitigates the effects on oocyte growth and ovulation, which is explained by a partial rescue of oocyte-somatic cell communication and folliculogenesis. Together, this work demonstrates that mitochondrial function influences communication of oocyte with follicular somatic cells and suggests that the balanced expression of modulators of mitochondrial dynamics is critical for proper oocyte development.

## INTRODUCTION

Mitochondria play a key role in oocyte development (Chiaratti et al., 2018), termed oogenesis. The biogenesis of these organelles is heavily increased during oocyte growth (Jansen and De Boer, 1998) and likely accounts for the energetic needs of the oocyte and the early embryo (Collado-Fernandez et al., 2012). Furthermore, redistribution of mitochondria through the oocyte cytoplasm is fundamental to supply the energetic demands of key events such as meiotic progression and chromosome segregation (Coticchio et al., 2014; Udagawa et al., 2014; Wakai et al., 2014; Yu et al., 2010). In support of this, mitochondrial abnormalities in oocytes are tightly associated with disrupted meiotic spindle, embryonic developmental arrest, and sterility in humans and animals (Van Blerkom et al., 1995; Fragouli and Wells, 2015; Johnson et al., 2007; Reynier et al., 2001; Santos et al., 2006; Wai et al., 2010).

Interestingly, in oocytes mitochondria are more numerous, smaller, and rounder in appearance than in somatic cells (Chiaratti et al., 2018; Motta et al., 2000; Wassarman and Josefowicz, 1978). These morphological characteristics are determined by mitochondrial dynamics, the opposing processes of fusion and fission that regulate organelle activity, transport and degradation (Mishra and Chan, 2014; Schrepfer and Scorrano, 2016); cells lacking mitochondrial fusion have a severe defect in respiratory capacity and mitochondrial DNA (mtDNA) instability (Chen et al., 2003, 2005, 2010). Mitochondrial fusion is orchestrated by two dynamin-related GTPases - Mitofusins 1 (MFN1) and 2 (MFN2) on the mitochondrial outer membrane (Mishra and Chan, 2014; Schrepfer and Scorrano, 2016).

Given the unique architecture of the mitochondrial network in oocytes, it is likely that *Mfn1* and *Mfn2* are more tightly regulated in the female germline (Liu et al., 2016a; Wakai et al., 2014; Zhang et al., 2016; Zhao et al., 2015). Indeed, while the ablation of fission leads to mitochondrial elongation in oocytes (Udagawa et al., 2014), the overexpression of *Mfn1* and *Mfn2* does not lead to mitochondrial elongation (Wakai et al., 2014), suggesting maintaining a fragmented population is critical for the oocyte. Moreover, mice with oocyte-specific deficiency for mitochondrial fission are infertile (Udagawa et al., 2014), whereas overexpression of fusion genes causes marked mitochondrial aggregation and damages oocyte viability (Wakai et al., 2014). Although there is a clear link between mitochondrial dynamics and oocyte fertility (Liu et al., 2016b; Udagawa et al., 2014; Wakai et al., 2014; Yu et al., 2010; Zhang et al., 2016), many questions remain: why do mitochondrial populations in the female gamete have such a unique morphology relative to somatic cells? How do these changes in shape relate to the role of mitochondria in oocyte development? Is the essential role of pro-fusion proteins MFN1 and MFN2 in early embryogenesis (Chen et al., 2003) linked to oocyte development? To tackle these questions and specifically address the role of mitofusins in the female gamete, we generated oocyte-specific conditional knockouts (cKO) of *Mfn1, Mfn2,* and both *Mfn1* & *Mfn2* (*Mfn1&2*) in mice. We found that the deletion of *Mfn1* alone, but not *Mfn2*, resulted in a complete loss of oocyte growth and a block in folliculogenesis at the preantral-to-antral follicle transition. Further characterization of *Mfn1*-cKO oocytes revealed impaired communication between the oocyte and its companion somatic cells. Unexpectedly, these defects were mitigated in mice with double loss of *Mfn1* and *Mfn2* in oocytes, which was explained by a partial rescue of oocyte-somatic cell communication and folliculogenesis. These results demonstrate that *Mfn1,* but not *Mfn2*, is required for oocyte formation and ovulation, and that the balanced expression of *Mfn1* and *Mfn2* is a key determinant for oocyte-follicular cell communication.

## RESULTS AND DISCUSSION

### Mfn1 deletion impairs oocyte development, which is partially rescued by the double loss of Mfn1 and Mfn2

Oocyte development relies on i) a period of growth in which prophase I-arrested oocytes amass essential molecules and organelles needed later during oogenesis and embryogenesis; and, ii) a period of maturation characterized by the breakdown of the nuclear membrane – known as germinal vesicle (GV) – and progression of meiosis to the metaphase-II stage (Clarke, 2017). These stages of oocyte development take place inside ovarian follicles as somatic cells (e.g., granulosa and cumulus cells) provide an adequate environment for the oocyte until its ovation (Su et al., 2009).

To address the role of MFN1 and MFN2 in oocyte development, *Mfn1*^loxP^ and/or *Mfn2*^loxP^ males (Chen et al., 2007, 2010) were crossed to Zp3-Cre females to achieve homozygous deletion of these genes specifically in growing oocytes (de Vries et al., 2000). Conditional knockout females were first verified for the oocyte-specific loss of *Mfn1* and/or *Mfn2*. As expected, these transcripts were absent in oocytes (Figure 1A), but not in the surrounding cumulus cells (Figure S1A). Moreover, mitochondrial architecture in cumulus cells remained unaltered across groups (Figure S1B). Next, we assessed the impact of the knockouts on fertility by mating these females with WT males. Surprisingly, the deletion of *Mfn1* led to sterility in *Mfn1*-cKO and *Mfn1&2*-cKO females (Figure 1B), despite normal mating behavior and vaginal plug formation. In contrast, the number of pups born to *Mfn2*-cKO females was not affected (Figure 1B). In addition, females with heterozygous knockout oocytes (Zp3-Cre:*Mfn1*^+/−^, Zp3-Cre:*Mfn2*^+/−^, Zp3-Cre:*Mfn1*^+/loxP^ and Zp3-Cre:*Mfn2*^+/loxP^) demonstrated full viability and fertility, and pups were born at the expected (de Vries et al., 2000) Mendelian frequencies (data not shown).

**Figure 1.**
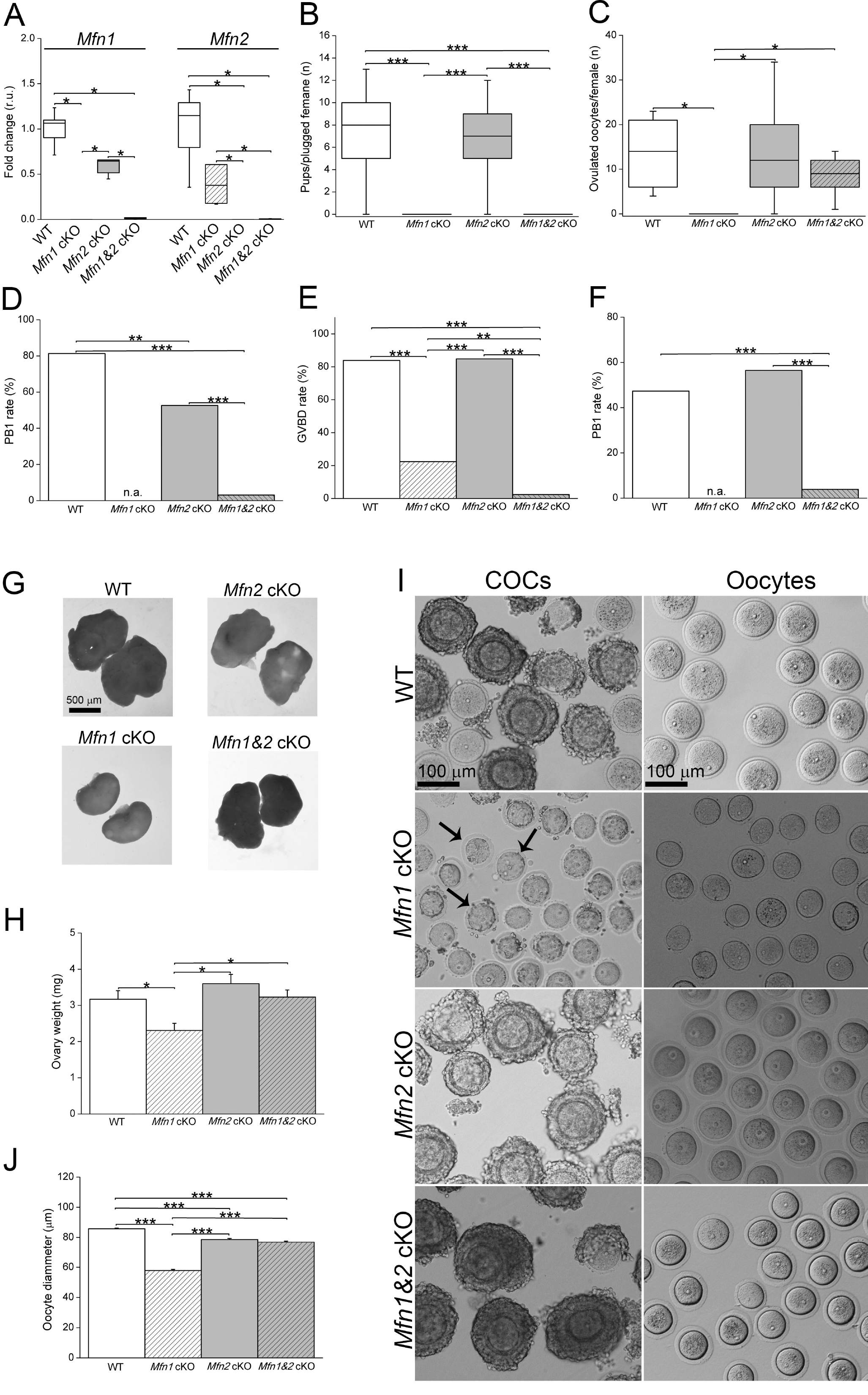
Mfn1 deletion impairs oocyte development, which is partially rescued by the double loss of Mfn1 and Mfn2. (A) Immature germinal-vesicle (GV) oocytes from 5-week-old mice were evaluated by qPCR to confirm the oocyte-specific loss of *Mfn1* and *Mfn2*. n = 5 samples/group. (B) Fertility test assay showing number of pups per plugged female. Note that deletion of *Mfn1* and *Mfn1&2* led to sterility. n = 20-52 matings/group. (C and D) Number of ovulated oocytes retrieved from oviducts of eCG- and hCG-primed mice (C). Rate of first polar body (PB1) in ovulated oocytes (D). (E and F) Rate of GV break down (GVBD; E) and PB1 extrusion (F) in oocytes matured in vitro. (G and H) Ovary morphology (G) and weight (H) in 5-week-old mice. n = 10-12 ovaries/group. (I) Cumulus-Oocyte Complexes (COCs) at the GV stage were retrieved from ovaries of eCG-primed mice aged 5 weeks (left). COCs were denuded of cumulus cells to denote oocytes (right). Note that most *Mfn1*-cKO COCs completely lacked surrounding cumulus cells (arrows), whereas grossly normal COCs were obtained from WT, *Mfn2*-cKO and *Mfn1&2*-cKO mice. (J) Diameter of immature GV oocytes from 5-week-old mice. Note the smaller size of *Mfn1*-cKO oocytes. n = 74-178 oocytes/group. In panels A, B and C, median (line inside the box), upper and lower quartiles (delimited by the box) and 1.5 times the interquartile range (whiskers). In panels H and J, mean ± SEM. n.a. = not analyzed; r.u. = relative units to the WT control. *P < 0.05. **P < 0.01. ***P < 0.0001. See also Figure S1.

Given that *Mfn1* is not required in the mouse proper for normal development (Chen et al., 2007), we considered that females lacking *Mfn1* and *Mfn1&2* were infertile due to an MFN1 deficiency in oocytes. In line with this, no ovulated oocytes were recovered from *Mfn1*-cKO mice (Figure 1C). However, the deletion of both *Mfn1* and *Mfn2* recovered the loss of ovulation: ~9 oocytes were recovered per *Mfn1&2*-cKO mice in comparison with ~14 oocytes from WT mice (Figure 1C).

To understand the reason for this difference, we assessed in ovulated oocytes the rate of first polar body (PB1) extrusion, a readout for meiotic progression to the metaphase-II stage. Only 3.1% of the oocytes that were ovulated by *Mfn1&2-cKO* mice contained PB1 (Figure 1D), while 61.4% of them were arrested at the GV stage. This finding was confirmed by in vitro maturation of GV oocytes (Figures 1E and 1F), indicating that *Mfn1&2*-cKO females were infertile due to ovulation of unviable oocytes. Mating of superovulated *Mfn1&2*-cKO females with WT males also confirmed this as the ovulated oocytes were not fertilized (data not show). The consequence of *Mfn1* ablation in the oocyte also extended to ovary development; in *Mfn* 1-cKO mice the ovary was ~62% smaller relative to WT, and ~60% relative to *Mfn1&2* cKO (Figures 1F and 1H). Moreover, immature GV oocytes retrieved from *Mfn1*-cKO ovaries were smaller and lacked companion cumulus cells when compared with *Mfn1&2*-cKO oocytes (Figures 1I and 1J).

Together, these data establish that *Mfn1* is essential for oocyte growth and ovulation, and that this essentiality is diminished in the absence of *Mfn2*.

### Folliculogenesis is blocked at the preantral-to-antral follicle transition in *Mfn1*-null oocytes

We hypothesized that failed ovulation (Figure 1C) and the smaller size of *Mfn1*-cKO ovaries and oocytes (Figures 1H and 1J) were a consequence of defective folliculogenesis (Udagawa et al., 2014). With activation of oocyte development, follicular cells start to differentiate and proliferate rapidly following a well-established pattern of development (Figure 2A) (Clarke, 2017). To address our hypothesis, ovaries were evaluated by histology and their follicles classified according to developmental stage as types 2, 3a, 3b, 4, 5a and 5b and 6 (Pedersen and Peters, 1968). In comparison with the WT group, *Mfn1*-cKO ovaries harbored a greater number of preantral follicles (types 2, 3a, 3b and 5a; Figure 2B). The number of antral type 6 follicles, however, was not increased in *Mfn1*-cKO ovaries (Figure 2B); instead, these were smaller and less developed than most antral follicles in WT ovaries (Figure 2C). This impact on antral follicle development is in agreement with recombination of loxP sites occurring in our system in follicles types 3b and 4 (de Vries et al., 2000). Key steps of folliculogenesis following these stages are the massive proliferation of granulosa cells and the formation of an antrum that subdivides granulosa cells into two populations: the mural granulosa and the cumulus cells (Figure 2A) (Clarke, 2017).

**Figure 2.**
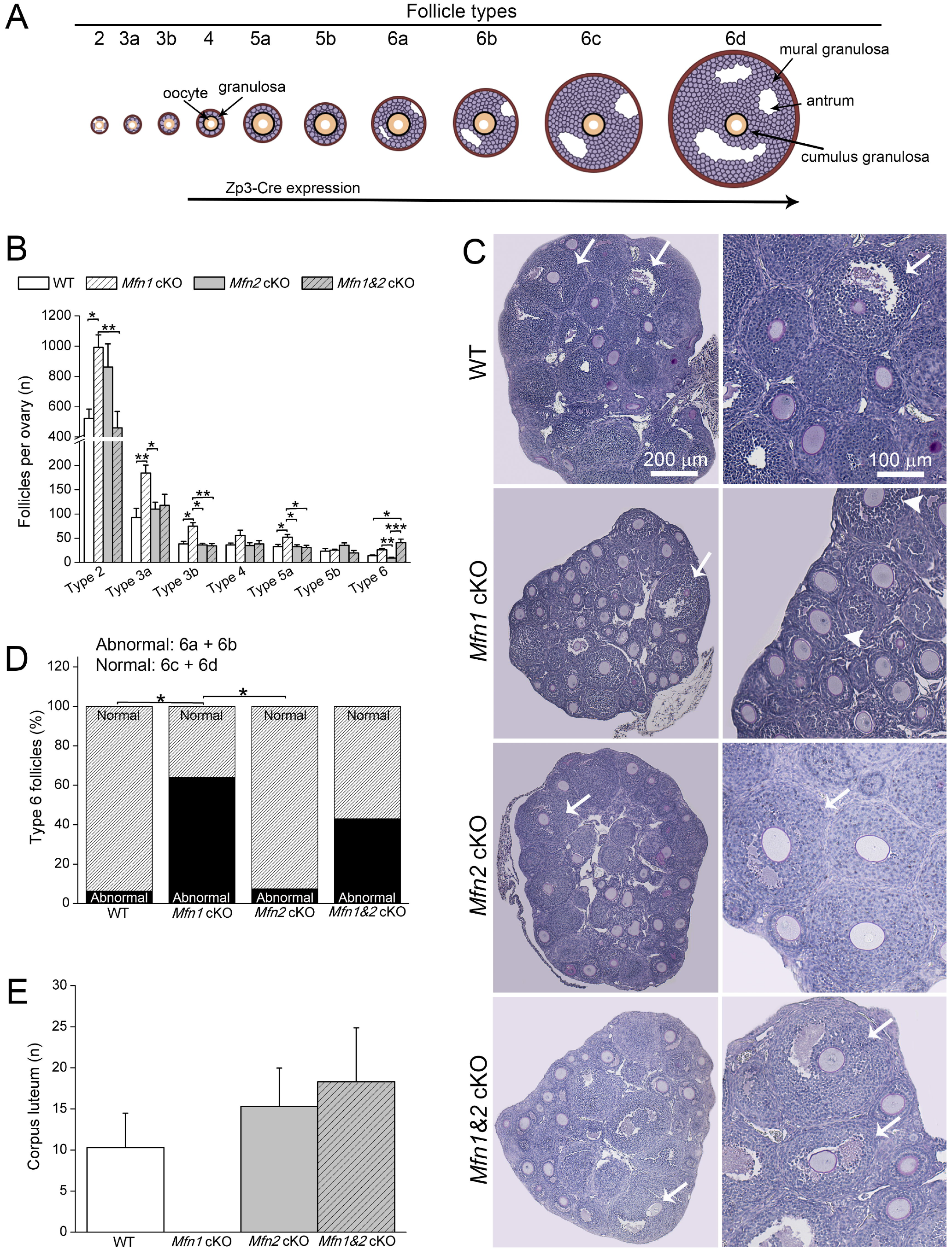
Folliculogenesis is blocked at the preantral-to-antral follicle transition in Mfn1-cKO oocytes. (A) Cartoon illustrating follicle development as proposed by Pedersen and Peters (1968). Cre recombinase is expressed in Zp3-Cre mice starting in follicle types 3b and 4. (B-D) Histological evaluation of ovaries from 3-week-old mice and follicular classification as types 2, 3a, 3b, 4, 5a, 5b and 6 (B). Follicles classified as type 6 represent the sum of types 6a, 6b, 6c and 6d. Types 1, 7 and 8 were not considered as these were mostly absent in 3-week-old mice. Representative histological micrographs of ovaries (C). Arrows and arrowheads indicate, respectively, normal and abnormal type 6 follicles. Percentage of abnormal (types 6a and 6b - black bar) and normal (types 6c and 6d - striped bar) follicles (D). n = 6 ovaries/group. (E) Number of corpus luteum determined by histological analysis of ovaries from 8-week-old mice. n = 3-4 mice/group. In panels B and E, mean ± SEM. In panel D, median (line inside the box), upper and lower quartiles (delimited by the box) and 1.5 times the interquartile range (whiskers). r.u. = relative units to the WT control. *P < 0.05. **P < 0.01. ***P < 0.0001. See also Figure S2.

To further characterize the differences in antral follicles between WT and *Mfn1*-cKO mice, we subdivided type 6 follicles into four subcategories: 6a, 6b, 6c and 6d (Figures 2A and S2A), ranging from poorly-developed (6a and 6b - abnormal) to well-developed antral follicles (6c and 6d - normal). In WT mice, a low percentage of follicles was classified as abnormal (Figures 2D, S2B and S2C), while in *Mfn1*-cKO mice 64.0% were classified as abnormal (Figures 2D, S2B and S2C). Interestingly, the *Mfn1&2*-cKO mice presented an intermediate phenotype with 43.0% classified as abnormal (Figures 2D, S2B and S2C).

Following ovulation, residual follicular cells differentiate into a gland, termed corpus luteum, that is mainly involved with progesterone production during early pregnancy. Consistent with a failure of *Mfn1*-cKO follicles to ovulate, *Mfn1*-cKO, but not *Mfn1&2*-cKO, ovaries completely lacked corpus luteum relative to the WT control (Figure 2E). Together, these results show that the loss of *Mfn1* in oocytes results in a block at the preantral-to-antral follicle transition, and that the combined loss of *Mfn1* and *Mfn2* mitigates this block in follicular development.

### Impaired folliculogenesis in Mfn1-deficient oocytes is associated with abnormal mitochondrial architecture and function

MFN1 and MFN2 are key regulators of mitochondrial health (Chen et al., 2003, 2005, 2007, 2010; Mishra and Chan, 2014; Papanicolaou et al., 2012; Schneeberger et al., 2013; Schrepfer and Scorrano, 2016; Sebastian et al., 2012), which in turn is linked with oocyte development (Van Blerkom et al., 1995; Fragouli and Wells, 2015; Johnson et al., 2007; Reynier et al., 2001; Santos et al., 2006). Given the differences observed, we hypothesized there might be differences specific to the loss of *Mfn1* that were mitigated by the double loss of *Mfn1* and *Mfn2*. Indeed, ultrastructure analysis of *Mfn1*-cKO oocytes revealed pronounced defects in mitochondria including: a) a greater number of less electron dense organelles; b) a decreased number of cristae; and, c) an increased vesiculation of the inner membrane (Figures 3A-3C and S3A). These mitochondrial abnormalities were present, albeit, to a lower extent in *Mfn1&2*-cKO than in *Mfn1*-cKO oocytes (Figures 3A-3C and S3A), suggesting that defective mitochondrial architecture is linked with the block of *Mfn1* cKO on follicle development.

**Figure 3.**
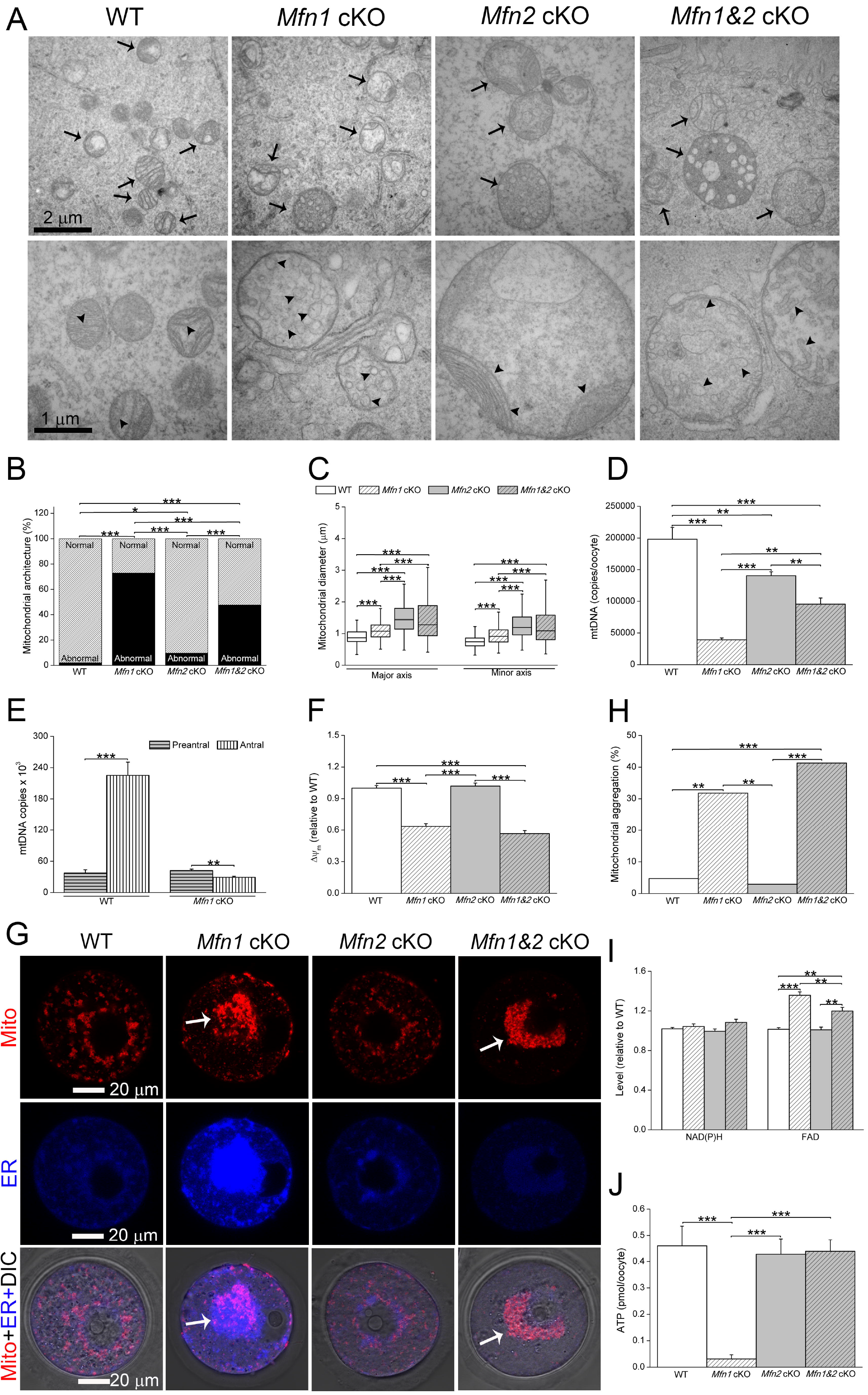
Impaired folliculogenesis in Mfn1-deficient oocytes is associated with abnormal mitochondrial architecture and function. (A) Representative transmission electron microscopy micrographs of oocytes. Note that mitochondrial morphology is abnormal in *Mfn1*-cKO oocytes with evidence of organelle swelling, vesiculation of the inner membrane, decreased number of cristae and lighter matrix. Arrows indicate mitochondria whereas arrowheads denote inner mitochondrial membranes and internal vesicles. (B) Percentage of mitochondria with normal (striped bar) and abnormal (black bar) ultrastructure characteristics in oocytes. Note that *Mfn1*-cKO oocytes depict an increased incidence of mitochondria with abnormalities. n = 53-78 samples/group. (C) Major and minor mitochondrial diameters in oocytes. n = 173-415 samples/group. (D and E) Number of mtDNA copies in immature germinal-vesicle (GV) oocytes from 5-week-old mice primed with eCG. Note that *Mfn1* loss impaired mtDNA accumulation more sharply than *Mfn1&2* cKO (D). Number of mtDNA copies in oocytes from preantral (12-day-old mice) and antral (32-day-old mice) follicles (E). (F) Relative mitochondrial membrane potential (ΔΨm) in immature GV oocytes estimated using TMRE. n = 89-229 samples/group. (G and H) Representative confocal microscopy micrographs (G) of immature GV oocytes stained with MitoTracker CMXRos (mitochondria - Mito) and ERTracker (endoplasmic reticulum - ER). ER staining is presented as a cytoplasmic reference. Percentage of GV oocytes with mitochondrial aggregation (H). Note the higher level of mitochondrial aggregation in *Mfn1*-cKO and *Mfn1&2*-cKO oocytes (arrows). DIC = differential interference contrast. (I) Relative levels of NAD(P)H and FAD in GV oocytes. n = 289-794 samples/group. (J) ATP content in immature GV oocytes. n = 15-21 samples/group. In panels C, median (line inside the box), upper and lower quartiles (delimited by the box) and 1.5 times the interquartile range (whiskers). In panels D, E, F, I and J, mean ± SEM. *P < 0.05. **P < 0.01. ***P < 0.0001. See also Figure S3.

Deficient fusion has been shown to lead to alterations in nucleoid heterogeneity and mtDNA instability and depletion (Chen et al., 2003, 2005, 2007, 2010). In turn, mtDNA copy number has been linked with oocyte quality in mice and humans (Fragouli and Wells, 2015; Reynier et al., 2001; Santos et al., 2006; Wai et al., 2010). We therefore next assessed levels of mtDNA, the regulation of which is intimately tied with mitochondrial function. To investigate whether the folliculogenesis block in *Mfn1*-cKO oocytes was associated with changes in mtDNA stability, we measured mtDNA copy number in immature GV oocytes. *Mfn1*-cKO oocytes had a ~5-fold reduction in mtDNA copy number relative to the WT control (Figure 3D). In addition, a comparison of *Mfn1*-cKO oocytes from preantral and antral follicles revealed that the content of mtDNA dropped with follicular development (Figures 3E and S3B). The loss of mtDNA was mitigated in *Mfn1&2*-cKO oocytes which had only a mild depletion (~2-fold reduction) relative to the WT control (Figure 3D). These data support a key role of *Mfn1* for mtDNA synthesis and/or stability in oocytes. Furthermore, our findings demonstrate that the double deletion of *Mfn1* and *Mfn2* mitigates the loss of mtDNA in oocytes. To further characterize the impact of *Mfn1* deletion on mitochondrial physiology and identify defects specific to the loss of Mfn1, but not *Mfn1&2*, we stained GV oocytes with TMRE to assess mitochondrial membrane potential (ΔΨm). Consistent with the observed defects in mitochondrial architecture, oocytes lacking *Mfn1* and *Mfn1&2* displayed a similar drop of ~40% in the ΔΨm (Figures 3F). Additionally, a large percentage of these oocytes presented mitochondrial aggregates (Figures 3G, 3H and S3C). However, the content of oxidized flavin adenine dinucleotide (FAD) was less altered in *Mfn1&2*-cKO oocytes in comparison with those lacking only *Mfn1* (Figures 3I). Moreover, *Mfn1&2*-cKO and WT oocytes contained similar levels of ATP, which was decreased by ~10 fold in the *Mfn1*-cKO group (Figure 3J). Overall, the mitochondrial defects were milder in *Mfn1*-cKO oocytes, indicating that the rescue effect of combined loss of *Mfn1* and *Mfn2* extended to mitochondrial health. These results link abnormal mitochondrial architecture, mtDNA depletion, increased levels of FAD, and lower levels of ATP with impaired folliculogenesis in *Mfn1*-cKO oocytes.

### The loss of Mfn1 dysregulates oocyte-somatic cell communication

The evident loss of cumulus cells only in oocytes with *Mfn1* single deletion (Figure 1I) hinted at a role for balanced expression of MFN1 and MFN2 in proliferation of follicular somatic cells. This hypothesis is in agreement with a milder effect of *Mfn1&2* cKO on antral follicle development and ATP storage in oocytes (Figures 2D and 3J). Moreover, *Mfn1*-cKO ovaries presented a more altered pattern of expression (Figures 4A and 4B) of genes implicated in granulosa cell proliferation and follicle development (Knight and Glister, 2006). The interaction of the oocyte with its companion somatic cells is mediated by oocyte-derived factors that stimulate proliferation and differentiation of these cells (Clarke, 2017). This communication is critical for oocyte development as, among other roles, companion somatic cells physically and metabolically interact with the oocyte to supply it with energetic molecules including ATP (Su et al., 2009; Sugiura et al., 2007). To provide additional evidence that the single *Mfn1* deletion impaired cellular proliferation, mice were injected for 2 h with BrdU and assessed as for the percentage of BrdU-positive follicular cells. Accordingly, BrdU incorporation was more sharply impaired by loss of *Mfn1* than the dual loss of *Mfn1* and *Mfn2* (Figures 4C and S4A).

**Figure 4.**
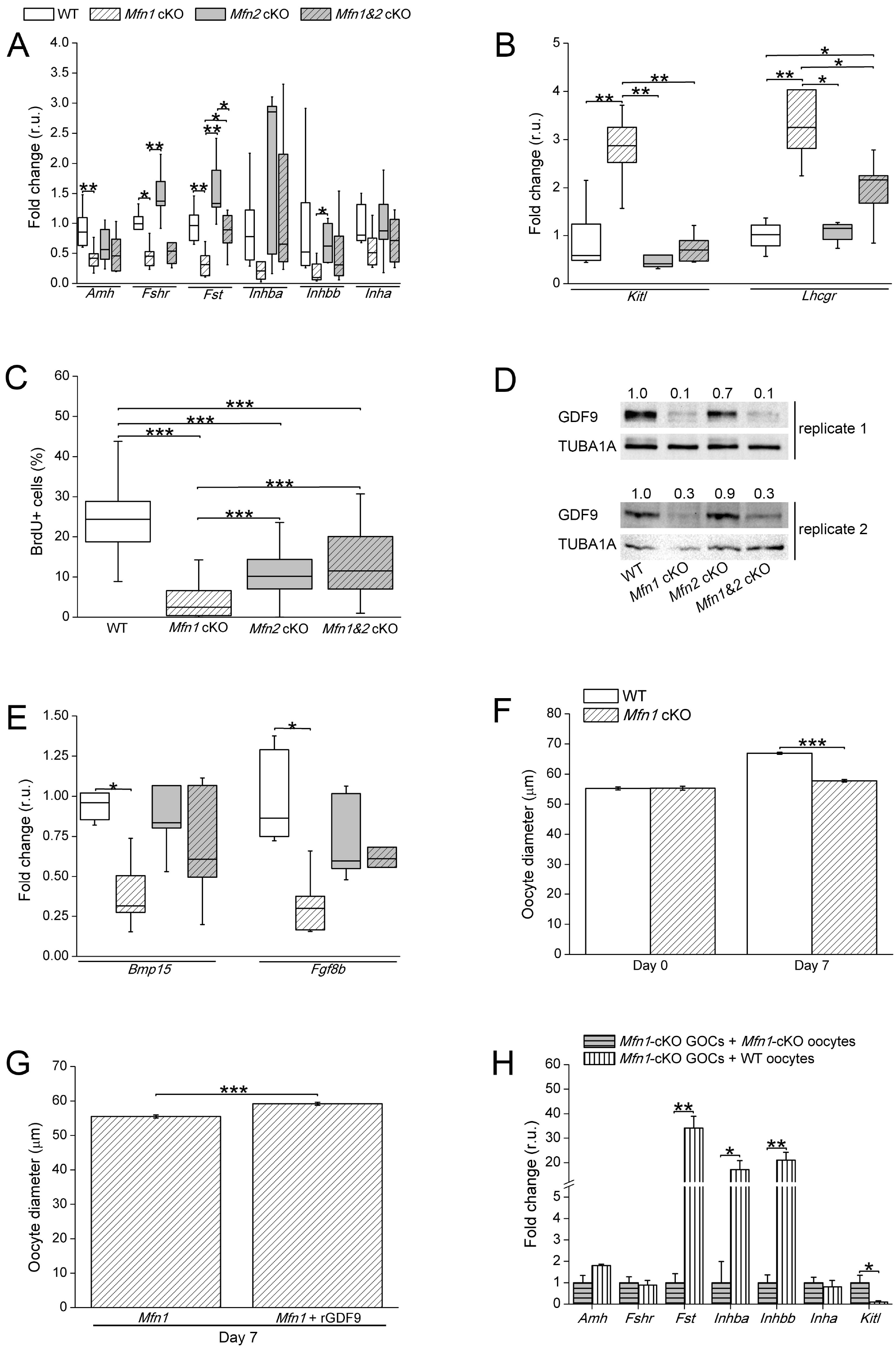
The loss of Mfn1 dysregulates oocyte-somatic cell communication. (A and B) Five-week-old ovaries were used to determine the levels of transcripts positively (A) and negatively (B) associated with follicle development. n = 8 ovaries/group. (C) Percentage of follicle cells positive for BrdU staining in ovaries from 8-week-old mice. (D) Evaluation by immunoblot of the level of GDF9 in germinal-vesicle (GV) oocytes. TUBA1A was used as sample input. Values above blots indicate the level of GDF9 normalized by TUBA1A (in relation to the WT control). (E) Transcriptional levels of *Bmp15* and *Fgf8b* in immature GV oocytes from 5-week-old mice. n = 5 samples/group. (F and G) Oocyte diameter before (n = 18-21/group) and after (n = 58-84/group) culture of Granulosa-Oocyte Complexes (GOCs) in vitro (F). Oocyte diameter after culture of *Mfn1*-cKO GOCs with (n = 43) or without (n = 28) recombinant GDF9 (rGDF9) (G). (H) Coculture of *Mfn1*-cKO GOCs with WT and *Mfn1*-cKO GV oocytes (freed of cumulus cells) from 5-week-old mice. After culture, GOCs were used for evaluation of gene expression. In panels A, B, C and E, median (line inside the box), upper and lower quartiles (delimited by the box) and 1.5 times the interquartile range (whiskers). In panels F, G and H, mean ± SEM. *P < 0.05. **P < 0.01. ***P < 0.0001. r.u. = relative units to the WT control. In (H), expressed in relation to *Mfn1*-cKO GOCs cocultured with *Mfn1*-cKO oocytes. See also Figure S4.

Next, we investigated whether impaired oocyte-somatic cell communication could explain the deficient proliferation of follicular cells in *Mfn1*-cKO ovaries. GDF9 is the most well-known oocyte-derived factor responsible for mediating interaction with follicular somatic cells (Clarke, 2017; Su et al., 2009). For instance, GDF9-deficient mice are infertile due to a block in folliculogenesis secondary to failed growth and differentiation of follicular somatic cells (Dong et al., 1996; Elvin et al., 1999). In accordance with our hypothesis, GDF9 was found to be downregulated in both *Mfn1*-cKO and *Mfn1&2*-cKO oocytes (Figures 4D and S4B). Additionally, *Mfn1*-cKO, but not *Mfn1&2-cKO*, oocytes expressed lower levels of *Bmp15* and *Fgf8b* in comparison with WT (Figure 4E). BMP15 and FGF8B are additional oocyte-derived factors involved mainly with the metabolic regulation of surrounding somatic cells (Su et al., 2009; Sugiura et al., 2007). Therefore, these findings indicate that *Mfn1* cKO disrupts communication of the oocyte with follicular somatic cells.

To further confirm that impaired oocyte-somatic cell interaction prevented the growth of *Mfn1*-cKO oocytes, we took advantage of an in vitro system of Granulosa-Oocyte Complex (GOC) culture. In this system, GOCs are derived from preantral secondary follicles and kept in culture for 7 days to develop to a stage comparable with oocytes from antral follicles (O’Brien et al., 2003). First, we used this system to validate our in vivo findings (Figure 1J) that *Mfn1*-cKO oocytes are reduced in growth. Whereas WT oocytes grew from 55.2 to 66.9 μm, *Mfn1*-cKO oocytes grew only from 55.3 to 57.8 μm (Figure 4F). To determine if exogenous addition of GDF9 rescued the growth of *Mfn1*-cKO oocytes that lack GDF9 expression, we cultured GOCs in the presence of recombinant GDF9 (rGDF9). Supplementation with rGDF9 was sufficient to partially rescue growth in *Mfn1*-cKO oocytes (Figure 4G).

Finally, *Mfn1*-cKO GOC were co-cultured with either WT or *Mfn1*-cKO oocytes to confirm the inability of *Mfn1*-null oocytes to modulate granulosa cell gene expression (Sugiura et al., 2007; Udagawa et al., 2014). Indeed, co-culture of *Mfn1*-cKO GOCs with WT oocytes upregulated *Fst*, *Inhba* and *Inhbb*, and downregulated *Kitl* (Figure 4H). Importantly, *Fst* and *Kitl* were, respectively, downregulated and upregulated in *Mfn1*-cKO ovaries (Figures 4A and 4B), providing evidence that co-culture with WT oocytes retrieved an expression pattern compatible with follicle development (Knight and Glister, 2006; Udagawa et al., 2014). Together, these data support that single deletion of *Mfn1* in growing oocytes impairs signaling to surrounding somatic cells, thereby preventing oocyte growth and follicular cell development.

### Conclusions

Our present results support three main conclusions: a) the oocyte relies on *Mfn1*, but not *Mfn2*, for growth and ovulation; b) the effect of *Mfn1* deletion is explained by disruption of oocyte-somatic cell communication and follicular cell proliferation; and, c) loss of *Mfn2* alleviates the impact on intercellular communication and follicular cell proliferation, mitigating the defects of *Mfn1*-null oocytes.

Herein we investigate the role of pro-fusion proteins MFN1 and MFN2 in the female germline during a phase of massive oocyte growth, when essential molecules and organelles are amassed to support further oocyte and embryo development (Clarke, 2017; Collado-Fernandez et al., 2012). In agreement with the hypothesis of upregulated fission in oocytes (Chiaratti et al., 2018; Udagawa et al., 2014; Wakai et al., 2014), ablation of pro-fusion genes do not enhance mitochondrial fragmentation. Yet, oocytes do express both mitofusins and ablation of *Mfn1* heavily impairs mitochondrial architecture, function and transport. These present findings corroborate previous reports (Chen et al., 2005; Tang, 2015; Udagawa et al., 2014; Wakai et al., 2014; Zhang et al., 2016) and unmask a key role of MFN1 in oocytes in comparison with MFN2. Previous reports have shown that mitofusins operate redundantly and non-redundantly in several tissues including placenta, neurons, hepatocyte, cardiomyocytes and skeletal muscle (Chen et al., 2007, 2010; Dietrich et al., 2013; Papanicolaou et al., 2012; Schneeberger et al., 2013; Sebastian et al., 2012). In comparison, our present findings highlight the importance of distinct, non-redundant roles of MFN1 and MFN2 in the growing oocyte.

Rather than a direct consequence of mitochondrial loss of function, the impact of MFN1 deficiency in the oocyte is explained by disruption of oocyte-somatic cell communication. This conclusion is supported by previous reports showing cumulus cells provide the oocyte with energetic molecules (Su et al., 2009; Sugiura et al., 2007), enabling oocyte development even when mitochondria is dysfunctional (Collado-Fernandez et al., 2012; Johnson et al., 2007). Moreover, loss of both MFN1 and MFN2, which alleviates the impact on intercellular communication and follicular cell proliferation, mitigates the mitochondrial defects and enables oocyte growth and ovulation. Taking into account the mitohormesis theory (Yun and Finkel, 2014), these findings might be explained by a possible mitochondrial stress induced by loss of *Mfn1*, which in turn could have activated a signaling pathway leading to disrupted oocyte-somatic cell communication. However, the link between loss of *Mfn1* and disrupted oocyte-somatic cell communication remains an open question. In brief, this work reframes the role of mitochondria in oocyte biology by identifying a specific role for a mitochondrial protein in intercellular signaling during oocyte development. Also, it suggests that the balanced expression of *Mfn1* and *Mfn2* is determinant for the growing oocyte. Future work should dissect how mitochondria interfere with oocyte-somatic cell communication.

## Supporting information

## ACKNOWLEDGMENTS

We deeply thank Dr. Hugh J. Clarke (McGill University) and Dr. John J. Eppig (The Jackson Laboratory), for suggestions on the manuscript; Dr. Felipe R. Teixeira (DGE/UFSCar), Dr. Marta Giacomello (University of Padova), Dr. Nadja C. Souza-Pinto (IQ/USP), Ana P. Dezam, Katiane Tostes, Monique Maroldi, and Natally P. Pierri, for invaluable assistance with mice, experiments and analyzes; and, José A. Maulin and Maria D. Seabra Ferreira (LMME/FMRP/USP), for technical support with analysis by transmission electron microscopy. This work was funded by the São Paulo Research Foundation (FAPESP/Brazil – grants # 2009/54035-4, 2012/50231-6, 2017/05899-2, 2017/04372-0, 2016/11935-9, 2016/11942-5, 2016/07868-4 and 2018/06119-3) and the Coordenação de Aperfeiçoamento de Pessoal de Nível Superior (CAPES/Brazil – finance code 001). We apologize to colleagues whose work could not be cited owing to editorial policies.

## AUTHOR CONTRIBUTIONS

Conceptualization, F.V.M and M.R.C; Methodology, F.V.M., F.E.G.G., M.M.S., and M.R.C.; Investigation, K.F.C., B.M.G., T.S.M., A.F.Z., C.H.M., F.H.C.S, M.P.G., J.D.A.N., F.K.S.R., F.D.S., M.M.S., and M.R.C.; Resources, F.V.M., F.E.G.G., M.M.S., and M.R.C.; Writing – Original Draft, M.R.C.; Writing – Review & Editing, L.P. and M.R.C.; Supervision, M.M.S. and M.R.C.; Project Administration, M.R.C.; Funding Acquisition, M.R.C.

## EXPERIMENTAL PROCEDURES

### Mouse breeding and fertility test

All experiments were performed in compliance with the regulations and policies of the National Council for Control of Animal Experimentation (CONCEA, Brazil) and were approved by the Animal Care and Use Committee at Universidade de São Paulo (USP) and Universidade Federal de São Carlos (UFSCar). Mfn^loxP^ conditional mice (Chen et al., 2007) were obtained from the Mutant Mouse Regional Resource Centers (RRID: MMRRC Cat# 029901-UCD, RRID:MMRRC_029901-UCD and MMRRC Cat# 029902-UCD, RRID:MMRRC_029902-UCD) whereas Zp3-Cre transgenic mice (de Vries et al., 2000) were supplied by the Jackson Laboratory (RRID: IMSR Cat# JAX:003651, RRID:IMSR_JAX:003651). Before use, Zp3-Cre mice were mated with heteroplasmic C57BL/6N females containing mtDNA of NZB/BINJ and C57BL/6N origin (Machado et al., 2015).

Wild-type (Zp3-Cre:*Mfn1*^+/+^:Mfn2^+/+^), *Mfn1*-cKO (Zp3-Cre:*Mfn1*^loxP/−^;*Mfn2*^+/+^) and *Mfn2*-cKO (Zp3-Cre:*Mfn1*^+/+^:*Mfn2*^loxP/-^) mice were obtained by mating heteroplasmic Zp3-Cre females to *Mfn*^loxP/loxP^ males, followed by mating of Zp3-Cre:*Mfn*^+/loxP^ males to Zp3-Cre:*Mfn*^+/loxP^ females. *Mfn1&2*-cKO (Zp3-Cre:*Mfn1*^loxP/−^:*Mfn2*^loxP/−^) mice were similarly obtained, except that *Mfn1*^loxP/loxP^:*Mfn2*^loxP/loxP^ males were used. This breeding scheme ensures more effective recombination of loxP sites. Unless otherwise stated, all experiments were conducted using mice that aged 3-5 weeks.

Mice had fertility evaluated by mating 7-week-old females for up to one year with WT males. Only females with copulation plugs were considered during the fertility test.

### Mouse genotyping

Ear biopsies were digested for 3 h at 55°C using a solution containing 50 mM KCl, 10 mM Tris-Cl 2 mM MgCl_2_, 0.1 mg/ml gelatin, 0.45% Igepal CA-630, 0.45% Tween 20 and 100 μg/ml proteinase K (ThermoFisher Scientific). After digestion, samples were incubated at 95°C for 10 min, centrifuged at 10,000 x g for 5 min and the supernatant used for genotyping (Machado et al., 2015).

*Mfn1* and *Mfn2* were genotyped by PCR as suggested by the MMRRC, with few modifications. The presence of the Zp3-Cre transgene was determined by quantitative PCR (qPCR) based on the protocol provided by the Jackson Laboratory.

### Collection and in vitro culture of cells

Cumulus-Oocyte Complexes (COCs) and denuded oocytes were obtained as previously described (Nagy et al., 2003). Briefly, to obtain ovulated oocytes, females aged 3-5 weeks received an intraperitoneal injection of 5 I.U. of equine chorionic gonadotropin (eCG; Folligon, MSD Animal Health) and 5 I.U. of human chorionic gonadotropin (hCG; Chorulon, MSD Animal Health), given 46-47 h apart. Oviducts were removed from the mice 12-14 h after the hCG injection, and the ovulated oocytes retrieved in FHM medium (Nagy et al., 2003) by puncturing the ampulla with a 30G needle. These oocytes were denuded of surrounding cumulus cells by gentle pipetting in 0.3% hyaluronidase (Sigma-Aldrich) diluted in FHM (Nagy et al., 2003). Ovulated oocytes were then inspected for the presence of the first polar body (PB1) under a stereomicroscope (SMZ 745, Nikon).

To obtain COCs containing fully-grown immature oocytes at the germinal-vesicle (GV) stage, mice aged 3-5 weeks were injected with 5 I.U. of eCG and the ovaries removed after 44-46 h of injection. COCs were retrieved by puncturing the ovarian antral follicles in FHM medium (Nagy et al., 2003). Alternatively, Granulosa cell-Oocyte Complexes (GOCs) were obtained from 12-day-old females by incubating their ovaries in HEPES-buffered α-MEM (ThermoFisher Scientific) with 10 μg/ml type I collagenase and 10 μg/ml DNase for 30 min at 37°C (O’Brien et al., 2003). The ovary fragments were gently pipetted every 3 min to disrupt the tissue. COCs and GOCs had surrounding somatic cells removed by passing several times through a mouth-controlled glass pipette.

Immature GV oocytes were in vitro matured under mineral oil in 90-μl droplets of KSOM medium (Nagy et al., 2003) supplemented with 10% fetal bovine serum (ThermoFisher Scientific), 100 u/ml penicillin (ThermoFisher Scientific) and 100 u/ml streptomycin (ThermoFisher Scientific). A humidified incubator maintained at 37°C and containing 5% CO_2_ in air was used. To prevent GV breakdown (GVBD), oocytes were isolated in the presence of 100 μM IBMX (Sigma-Aldrich). Then, oocytes were cultured in IBMX-free medium and evaluated as for the GVBD (judged by absence of GV) and PB1 extrusion following 3 and 16 h of IBMX washout, respectively (Udagawa et al., 2014; Wakai et al., 2014).

The coculture experiment was performed as reported previously (Udagawa et al., 2014), with few modifications. Briefly, 10 GOCs were cultured under mineral oil for 24 h together with 15 denuded oocytes (from COCs) in 40-μl droplets of KSOM medium supplemented with 100 μM IBMX. A humidified incubator maintained at 37°C and containing 5% CO_2_ in air was used.

In vitro growth of oocytes was performed based on a previous report (O’Brien et al., 2003). Briefly, ~15 GOCs were cultured onto type I and III collagen 3.0-μm inserts (Corning) in 24-well plates with 750 μl of NaHCO_3_-buffered α-MEM supplemented with 1× ITS (Sigma-Aldrich), 10 μM cilostamide (Sigma-Aldrich), 10 mIU/ml FSH (Sigma-Aldrich) and, when applicable, 100 ng/ml of rGDF9 (R&D Systems). Two-thirds of the medium was replaced on the third day of culture. GOCs were cultured for 7 days in a humidified incubator maintained at 37°C and containing 5% CO_2_ in air.

When applicable, oocyte diameter was estimated under an inverted microscope (Ti-S, Nikon) using the NIS-Elements Basic Research software (Nikon).

### Histological and immunofluorescence analyses

Ovaries from 21- to 23-day-old females had the number of follicles evaluated as reported previously (Pedersen and Peters, 1968), with few modifications. Briefly, fixed ovaries were dehydrated, clarified, diaphonized, embedded in paraffin and sectioned at 3 μm thickness. Sections were stained with periodic acid and hematoxylin, and analyzed using an optical microscope. Follicles in every third section throughout the ovary were counted and classified as proposed by Perdersen and Peter (1968). Only follicles in which the section included the GV of the oocyte were considered. Ovaries from 8-week-old females were similarly processed to determine the number of corpus luteum. Follicular cell proliferation was assessed as previously reported (Udagawa et al., 2014), with few modifications. Briefly, 8-week-old females were injected with 3 mg BrdU (Sigma-Aldrich) 2 h prior to ovary collection. Ovaries were then fixed for 4 h in 4% paraformaldehyde, cryoprotected in a series of sucrose solutions (10-20%) in PBS at room temperature, embedded in OCT compound (Sakura Finetek) and frozen in cold acetone. Sectioned samples of 5-μm thickness were then thoroughly washed with PBS, fixed for 15 min in 2% paraformaldehyde and, incubated for 20 min with 10 mM sodium citrate and 0.05% Tween at 95°C. Unspecific staining was then blocked by incubation with 10% BSA in PBS for 30 min. Afterwards, slides were incubated overnight with mouse anti-BrdU (50-fold dilution; Sigma-Aldrich) primary antibody, thoroughly washed and incubated for 1 h with Alexa Fluor 594-tagged anti-mouse (800-fold dilution; ThermoFisher Scientific) secondary antibody. Cell nuclei were counterstained with 10 μg/ml DAPI (Sigma-Aldrich). The following wavelengths for excitation and emission, respectively, were considered: 591 nm and 614 nm (Alexa Fluor 594), and, 358 nm and 461 nm (DAPI). The percentage of replicating follicle cells (BrdU positive) in relation to DAPI-positive cells was determined by epifluorescence using the Ti-S microscope and the NIS-Elements Basic Research software. Only cells inside the follicle were considered during the analysis.

### Mitochondrial evaluation

Oocytes were incubated for 30 min at 37°C with 100 nM MitoTracker CMXRos (ThermoFisher Scientific) and 1 μM ERTracker (ThermoFisher Scientific) diluted in FHM for analysis of mitochondria and endoplasmic reticulum (ER) distribution, respectively. Similarly, oocytes were stained with 0.1 μM TMRE (ThermoFisher Scientific) in FHM for determination of mitochondrial membrane potential (ΔΨm). Oocytes treated with 10 μM CCCP for 30 min were used as negative control during ΔΨm evaluation. Oocytes were evaluated by confocal microscopy (LSM 780, Zeiss) considering the following wavelengths for excitation and emission, respectively: 579 nm and 599 nm (MitoTracker CMXRos), 374 nm and 430 nm (ERTracker), and, 485 nm and 535 nm (TMRE). Since the knockouts led to mitochondrial aggregation, which could potentially bias ΔΨm evaluation, oocytes were also analyzed by epifluorescence microscopy.

Oocyte autofluorescence, which is representative of the amount of oxidized flavin adenine dinucleotide (FAD) and reduced nicotinamide adenine dinucleotide (phosphate) (NAD[P]H), were determined using green (excitation = 465-495 nm; emission = 515-555 nm) and blue (excitation = 340-380 nm; emission = 435-485 nm) filters, respectively (Zeng et al., 2014).

Images were analyzed using the NIS-Elements Basic Research software or the ZEN lite (Zeiss). To control for inter-assay variation, oocytes analyzed in a specific experimental routine had fluorescence normalized by the mean fluorescence of WT oocytes analyzed in the same routine.

### ATP quantification

Oocytes used for ATP quantification were placed individually into 0.2 ml tubes in 1 μl PBS and kept at −80°C until use. Oocytes treated with 20 μM oligomycin for 1 h were used as negative control. The ATP content was measured based on the luciferin-luciferase reaction using the FLASC kit (Sigma-Aldrich), following manufacturer’s recommendations.

### Transmission electron microscopy

Ovaries used for transmission electron microscopy were obtained from 5-week-old mice after 44 h of eCG injection. These were fixed for 4 h at 4°C in 2% paraformaldehyde and 2% glutaraldehyde in 0.01 M cacodylate buffer (pH 7.3) with 3 mM CaCl. Following, ovaries were post-fixed for 1 h at 4°C with 2% osmium tetroxide and dehydrated in a grade series of ethanol. Samples were embedded in epoxy resin (Polysciences Inc.), cut into ultra-thin sections, mounted on 300-mesh copper grids and contrasted with uranyl acetate and lead citrate (Udagawa et al., 2014; Wakai et al., 2014). The sections were examined in a JEOL JEM - 100CXII transmission electron microscopy (Jeol Inc.). Images were analyzed using ImageJ (National Institutes of Health, Bethesda, MD).

### Evaluation of mtDNA copy number

The number of mtDNA copies in oocytes was determined by qPCR as described previously (Machado et al., 2015). Briefly, a 736-bp fragment encompassing nucleotides 3,455 to 4,190 of mtDNA was amplified using primers MT12 and MT13 (Table S1) and cloned into a pCR2.1-Topo-TA (ThermoFisher Scientific). Concentration of the resulting plasmid DNA was estimated based on UV spectroscopy and store at 0.2 × 10^9^ copies/μl at −80°C. This plasmid DNA was used to prepare a standard curve containing 0.2 × 10^7^, 0.2 × 10^6^, 0.2 × 10^5^, 0.2 × 10^4^ and 0.2 × 10^3^ copies/μl. A fresh dilution of this standard curve was always prepared before each qPCR reaction.

Individual oocytes collected in 1 μl of PBS with 0.01% BSA were digested for 3 h at 55°C in 4 μl of digestion solution (as described for mouse genotyping, except that 100 mg/ml proteinase K was used). After digestion, samples were incubated at 95°C for 10 min, centrifuged at 10,000 × g for 5 min and 5 μl of the supernatant diluted with 45 μl of ultrapure H_2_O (ThermoFisher Scientific).

The standard curve and samples were always amplified in parallel using the 7500 Fast Real-Time PCR System with the following cycling conditions: 1 cycle of 95°C for 10 min, 40 cycles of 95°C for 15 s and 62°C for 1 min. Each reaction consisted of 15 μl containing 200 nM of primers MT14 and MT15 (Table S1), 1x Power SYBR Green Master Mix and 5 μl of sample or standard-curve DNA. Samples were analyzed in triplicates and averaged for calculation of mtDNA copy number using the 7500 Fast Real-Time PCR Software (ThermoFisher Scientific).

### Evaluation of gene expression

Groups of 30 GV oocytes and 10 GOCs were placed into 0.2 ml tubes along with 10 μl PBS containing 1 U/μl RNase inhibitor (RNase OUT; ThermoFisher Scientific). Samples were snap frozen in liquid nitrogen and kept at −80°C until use. As reported previously (Macabelli et al., 2014), total RNA was extracted using TRIzol reagent (ThermoFisher Scientific), treated with DNase I (ThermoFisher Scientific) and reverse transcribed into cDNA using the High-Capacity cDNA Reverse Transcription kit (ThermoFisher Scientific). Gene transcripts were determined on the 7500 Fast Real-Time PCR as described previously (Macabelli et al., 2014), with few modifications. Briefly, each qPCR reaction consisted of 15 μl containing 1 μl of forward and reverse primers (200 nM each), 7.5 μl SYBR Green PCR Master Mix, 5 μl sample cDNA (20- to 77fold dilution) and 1.5 μl ultrapure H_2_O. A list of all primers used is presented at Table S1. These were designed using the PrimerBLAST software (National Institutes of Health) and confirmed regarding specificity of the amplified fragment by dissociation curve and gel electrophoresis.

All samples amplified for a specific target were run in duplicate in the same plate using the following cycling conditions: 95°C for 15 min followed by 45 cycles of 95°C for 15 s and 60°C for 1 min. Sample dilutions were determined based on standard curves generated for each gene-specific cDNA using five four-fold serial dilutions of cDNAs. These standard curves were also used for analysis of amplification efficiency (Livak and Schmittgen, 2001). Because all assays showed high amplification efficiency (~100%), data were linearized according to Livak and Schmittgen (Livak and Schmittgen, 2001). Samples that failed to amplify had cycle threshold-(CT) values regarded as equal to 45. The geometric average of CT values of two reference genes (*Ppia* and *Hprt*) was used to normalize gene expression data (Macabelli et al., 2014).

In case of ovaries, these were placed individually into cryo-tubes and snap frozen in liquid nitrogen. Ovaries were then macerated and subjected to total RNA extraction, followed by reverse transcription using 1 μg of total RNA and the High-Capacity cDNA Reverse Transcription kit. Gene transcripts were determined as described for oocytes, except that a 321-fold dilution of cDNA was used. Transcript (Table S1) amounts were evaluated as described above. Cumulus cells from groups of 30 oocytes were processed as described above, except that the PCR products were evaluated by electrophoresis on 2% agarose gel. DNA was then stained with SYBR Safe (ThermoFisher Scientific) and the signal detected with the Chemidoc XRS (BioRad).

### Immunoblot analysis

Groups of 100 denuded oocytes were placed into 0.2 ml tubes in 1 μl PBS, snap frozen in liquid nitrogen and kept at −80°C, until use. To compensate for the smaller diameter and, in turn, lower protein content, 200 oocytes were used in the case of *Mfn1* cKO. Samples were evaluated by western blot as previously (Wakai et al., 2014). Briefly, samples were lised, proteins size separated using 12% Bis-Tris Mini Gel and blotted onto PVDF membranes. Membranes were probed overnight with either rabbit anti-GDF9 (1000-fold dilution; Abcam) or mouse anti-TUBA1A (2000-fold dilution; ThermoFisher Scientific) primary antibodies, thoroughly washed and incubated for 1 h with HRP-conjugated secondary antibodies (10000-fold dilution; KPL). Then, membranes were treated with the Amersham ECL Select kit (GE) and the chemiluminescence signal detected with the Chemidoc XRS. For controlling sample input, protein levels were corrected by TUBA1A. Relative protein levels were determined using ImageJ.

### Statistical analysis

Statistical analysis was performed using the SAS University Edition v. 9.4 (SAS Institute Inc.). Data were tested for assumption of normal distribution and homogeneity of variance, and transformed when necessary. Experimental groups were then compared using unpaired t test (2 groups) or one-way ANOVA (4 groups), followed by Tukey post-hoc tests. Alternatively, groups were compared using a non-parametric test such as Wilcoxon Rank-Sum Test (2 groups) or Kruskal-Wallis Test (4 groups), followed by pairwise multiple comparison analysis. Binary variables were evaluated by binary logistic regression, followed by Tukey-Kramer post-hoc test. Correlation analysis was performed using Pearson’s Test. Differences with probabilities (P) < 0.05 were considered significant.

## SUPPLEMENTAL FIGURE LEGENDS

**Figure S1. Related to Figure 1. Mitofusins are not dysregulated in cumulus cells from Mfn-cKO oocytes.**

(A) Cumulus cells from WT, *Mfn2*-cKO and *Mfn1&2*-cKO oocytes had *Mfn1* and *Mfn2* amplified by PCR and run onto a 2%-agarose gel. *Ppia* was used as sample input. Because *Mfn1*-cKO oocytes mostly lacked cumulus cells (Figure 1I), these could not be used in this experiment. Five replicates were considered per experimental group.

(B) Representative transmission electron microscopy micrographs of oocytes and surrounding somatic cells. Arrows indicate oocyte mitochondria whereas arrowheads denote elongated mitochondria in somatic cells. Note that mitochondrial architecture in somatic cells is not different across groups.

**Figure S2. Related to Figure 2. *Mfn1* deletion arrests folliculogenesis at the preantral-to-antral follicle transition.**

(A) Representative histological micrographs denoting classification of early antral follicles, corresponding to type 6 (Pedersen and Peters, 1968), into four subcategories. Type 6a: ~128 cells and ~128 μm in diameter. Type 6b: ~186 cells and ~146 μm. Type 6c: ~338 cells and ~218 μm in diameter. Type 6d: ~483 cells and ~290 μm in diameter.

(B) Representative histological micrographs of Mfn-cKO ovaries from 21- to 23-day-old mice. Arrows and arrowheads indicate normal and abnormal type 6 follicles, respectively.

(C)Percentage of follicles classified as types 6a, 6b, 6c and 6d.

In panel C, median (line inside the box), upper and lower quartiles (delimited by the box) and 1.5 times the interquartile range (whiskers).

*P < 0.05.

**Figure S3. Related to Figure 3. *Mfn1* ablation results in oocytes with disrupted mitochondria.**

(A) Representative transmission electron microscopy micrographs oocytes. Note that mitochondrial morphology is abnormal in *Mfn1*-cKO oocytes with evidence of organelle swelling, vesiculation of the inner membrane and lighter matrix. Arrows indicate mitochondria whereas arrowheads denote inner mitochondrial membranes and internal vesicles.

(B) Relationship between mtDNA copy number and diameter in oocytes from 12-and 32-day-old mice. Both variables correlated (R) significantly in WT (R = 0.75; P < 0.0001), but not in *Mfn1*-cKO group (R = 0.21; P = 0.09).

(C) Representative confocal microscopy micrographs of immature germinal-vesicle (GV) oocytes stained with MitoTracker CMXRos (mitochondrial - Mito) and ERTracker (endoplasmic reticulum - ER). ER staining is presented as a cytoplasmic reference. DIC = differential interference contrast. Note the intense mitochondrial aggregation in *Mfn1&2*-cKO oocytes (arrows).

**Figure S4. Related to Figure 4. Oocyte-specific deletion of *Mfn1* impairs oocyte-somatic cell communication.**

(A) Representative fluorescence microscopy micrographs of BrdU-positive cells (arrowheads) in ovaries from 8-week-old mice. Ovary slides were counterstained with DAPI to determine cell nuclei. Dotted lines denote follicle area. Note the decreased number of BrdU-positive cells in *Mfn1*-cKO ovaries.

(B) Full-size images corresponding to blots presented on Figure 4D. Two different molecular weight markers were run on columns 1 and 2, whereas oocyte samples were run on columns 3-6. A total of 100 oocytes were used per group in each replicate, except for *Mfn1* cKO that 200 oocytes were used. Two biological replicates are show. Values above bands in column 1 and 2 represent molecular weights (kDa).

