## Supplementary figures and images for "Mitofusin 1 is required for the oocyte-granulosa cell communication that regulates oogenesis"

### Supplementary file 1

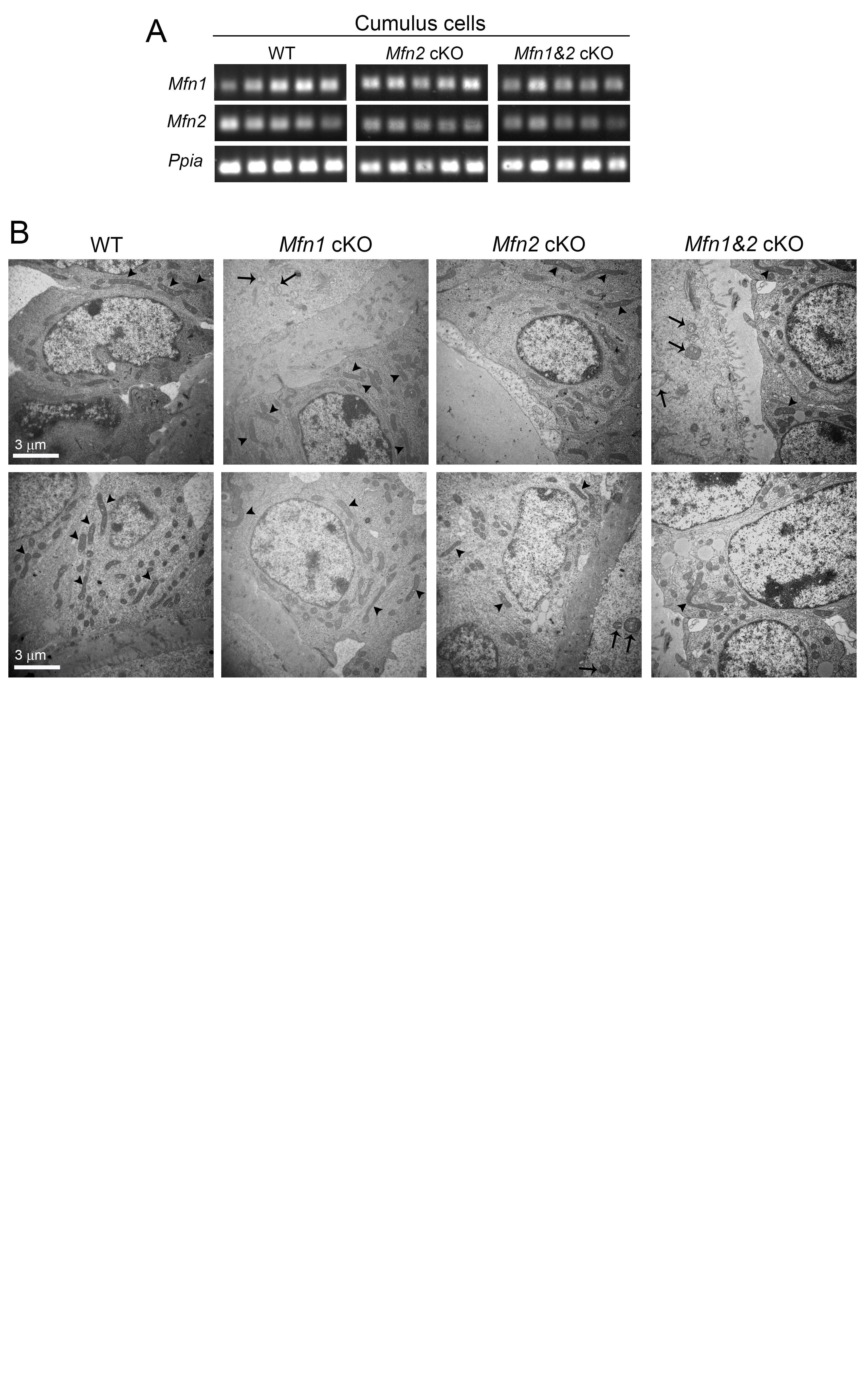

### Supplementary file 2

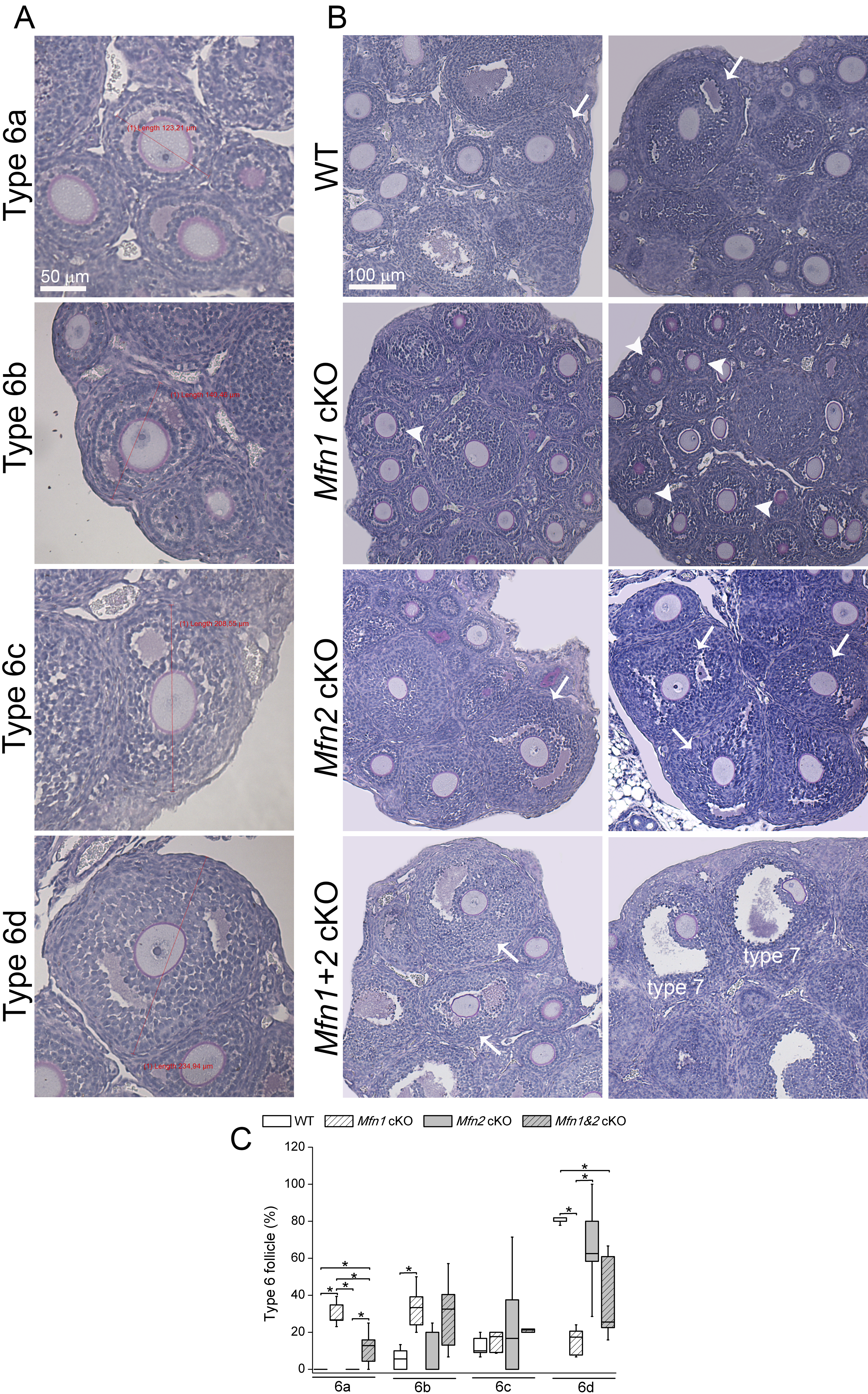

### Supplementary file 3

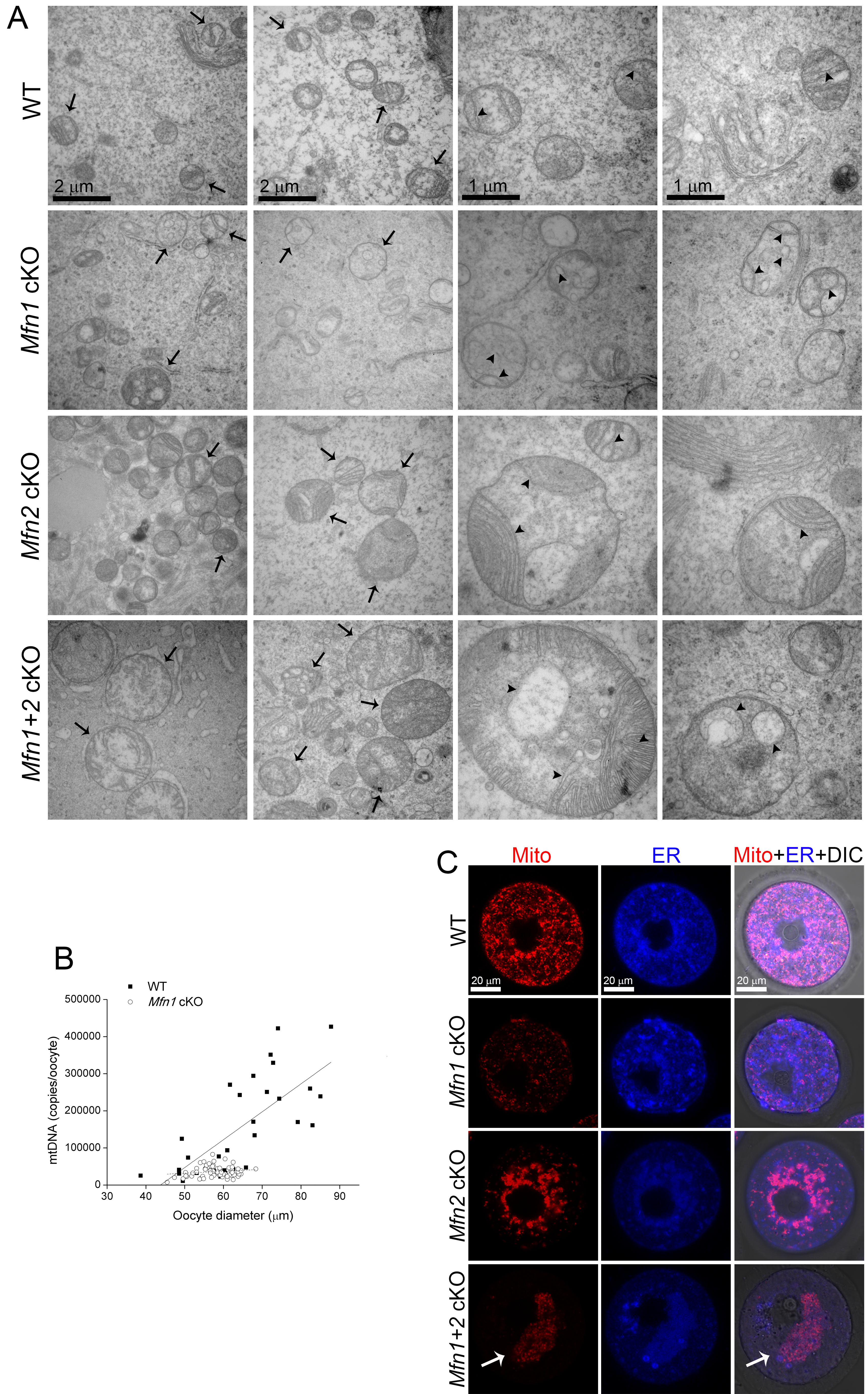

### Supplementary file 4

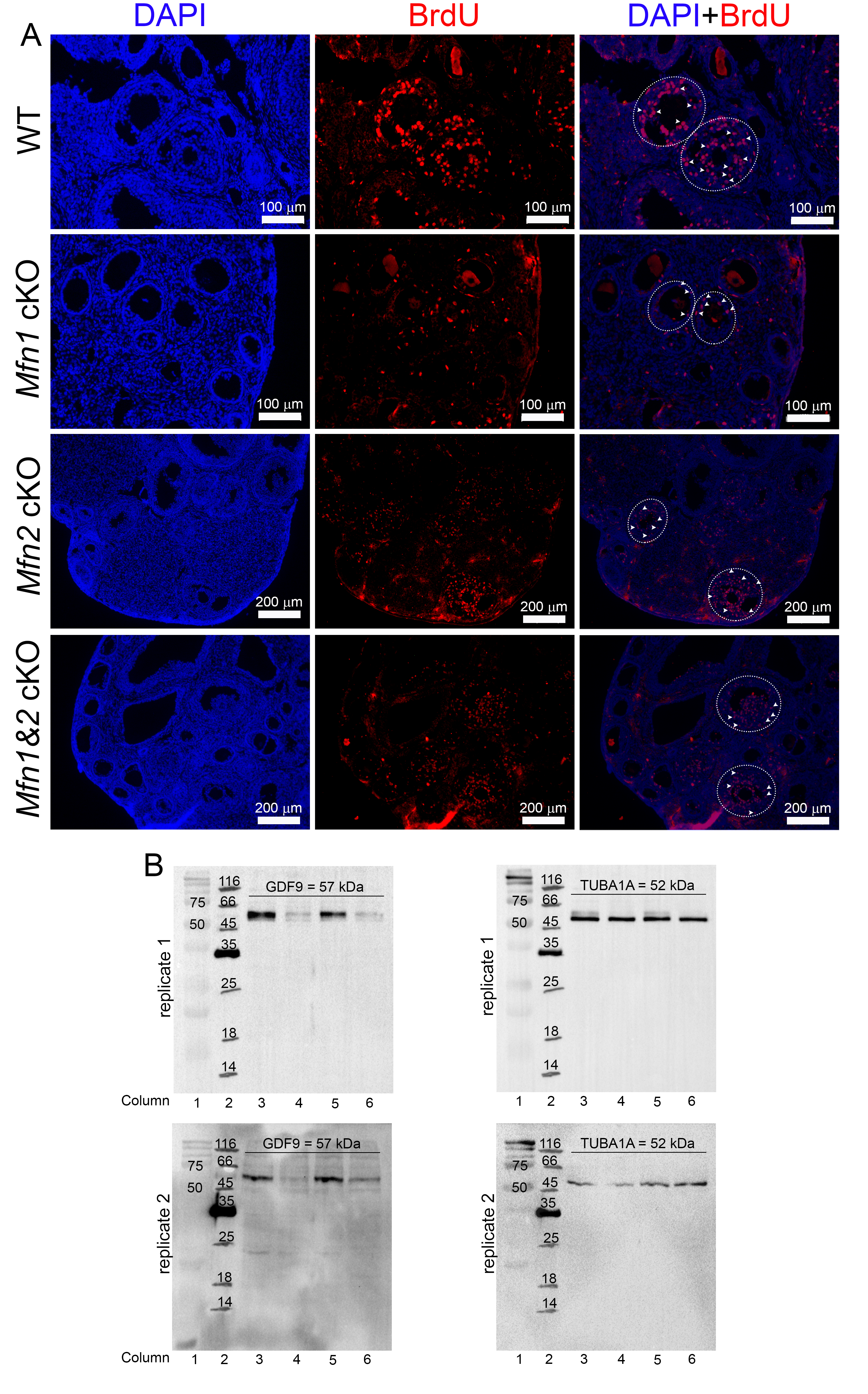
