## Supplementary material for "Mitofusin 1 is required for the oocyte-granulosa cell communication that regulates oogenesis"

**SUPLEMENTAL TEXT**

**Table S1. Primer sequences.**

| Primer name | Target gene (GenBank) | Sequence (5'-3') |
|---|---|---|
| MT14 | mtDNA (NC_005089.1) | CTCCGTGCTACCTAAACACCTTATC |
| MT15 | | GACCTAAGAAGATTGTGAAGTAGATGATG |
| MT12 | mtDNA (NC_005089.1) | CGCCCTAACAACTATTATCTTCC |
| MT13 | | GACCGTTTGTTTGTTGTTGAAA |
| Amh-F | *Amh* (NM_007445) | CTGGCTAGGGGAGACTGGAG |
| Amh-R | | TCGGGCTCCCATATCACTTC |
| Bmp15-F | *Bmp15* (NM_009757.5) | AAGGGAGAACCGCACGATTG |
| Bmp15-R | | TGTACATGCCAGGAACCTCTG |
| Fgf8b-F | *Fgf8* (NM_001166361.1) | GCTAATTGCCAAGAGCAACGG |
| Fgf8b-R | | AGCGCCGTGTAGTTGTTCTC |
| Fshr-F | *Fshr* (NM_013523.3) | GGAGGGCCAGGTCAACATACC |
| Fshr-R | | GAAGTCAGAGGTTTGCCGC |

| | | |
|---|---|---|
| Fst-F | *Fst* (NM_001301373.1) | TCATGGACCGAGGAGGATGT |
| Fst-R | | CCACGTTCTCACACGTTTCTTT |
| Hprt-F | *Hprt* (NM_013556) | GTTGGGCTTACCTCACTGCT |
| Hprt-R | | TCATCGCTAATCACGACGCT |
| Inha-F | *Inha* (NM_010564.4) | TATTCCGGCCATCCCAACAC |
| Inha-R | | CAGAAGATCTAGCAGGGGCG |
| Inhba-F | *Inhba* (NM_008380.1) | GGGACCCGAAAGAGAATTTGC |
| Inhba-R | | TCCTCTCAGCCAAAGCAAGG |
| Inhbb-F | *Inhbb* (NM_008381.3) | TTTGCAGAGACAGATGGCCT |
| Inhbb-R | | GAAGAAGTACAGGCGGACCC |
| Kitl-F | *Kitl* (NM_013598.2) | CACAGTGGCTGGTAACAGTTC |
| Kitl-R | | CCACACAAGGTCACGGGTAG |
| Lhcgr-F | *Lhcgr* (NM_013582.2) | CTCGCCCGACTATCTCTCAC |
| Lhcgr-R | | TTGAGGAGGTTGTCAAAGGCA |
| Mfn1-F | *Mfn1* (NM_024200.4) | CTGCTTCCTGAGTGTCGAGG |

| | | |
|---|---|---|
| Mfn1-R | | ATGCACAAGACAGCCAGCTT |
| Mfn2-F | *Mfn2* (NM_001285920.1) | CATGAGGCCTTCCTCCTCAC |
| Mfn2-R | | CAACTGCTCGTCCTGATGGA |
| Ppia-F | *Ppia* (NM_008907) | GTCTCCTTCGAGCTGTTTGC |
| Ppia-R | | GCGTGTAAAGTCACCACCCT |
